# Msc1 facilitates glucose starvation-induced remodeling of the nucleus-vacuole junction

**DOI:** 10.64898/2026.03.13.711511

**Authors:** Yusuke Mito, Shintaro Fujimoto, Saori Shinoda, Yasushi Tamura

## Abstract

The nucleus-vacuole junction (NVJ) is a membrane contact site between the nuclear envelope and the vacuole in yeast that undergoes dynamic remodeling in response to nutrient starvation. Here, we report that Msc1 is a glucose starvation (GS)-responsive NVJ factor. GS strongly induced Msc1 expression and promoted its accumulation at the NVJ. Although Msc1 is not essential for NVJ formation itself, loss of Msc1 impaired GS-dependent functional maturation of the NVJ, including stabilization and recruitment of multiple NVJ-associated proteins. Notably, GS-induced transcriptional activation of *NVJ1* was markedly attenuated in *msc1Δ* cells, suggesting that proper NVJ remodeling contributes to the execution of stress-responsive transcriptional programs. Together, these findings establish Msc1 as an upstream regulator linking GS to functional remodeling of the NVJ and associated transcriptional responses.

## Introduction

Eukaryotic cells dynamically reorganize organelle architecture and metabolic processes in response to cellular stress and nutrient conditions to maintain cellular homeostasis. Membrane contact sites (MCSs) are sites where the membranes of two organelles are closely apposed without fusion and contribute to such adaptive responses (Calì et al., 2025; Diokmetzidou and Scorrano, 2025; Prinz et al., 2020; Tamura et al., 2019; Voeltz et al., 2024). These structures undergo dynamic remodeling not only through changes in their molecular composition but also through alterations in their number, size, and spatial distribution (Elbaz-alon et al.; Hariri et al., 2018; Kohler and Büttner, 2021; Valm et al., 2017). For example, clusters of the ERMES complex that accumulate at mitochondria– ER contact sites (MERCS) dissociate under ER stress conditions, resulting in phospholipid accumulation within the ER membrane, ER expansion, and attenuation of stress (Kakimoto-Takeda et al., 2022). Another example the reciprocal regulation of ERMES-mediated MERCS and the mitochondria–vacuole contact site vCLAMP (vacuole and mitochondria patch) in response to carbon source availability, such that attenuation of one contact site is associated with expansion of the other (Elbaz-Alon et al., 2014; González Montoro et al., 2018; Hönscher et al., 2014). A recent study further demonstrates that distinct MCSs can be coordinately regulated and influence one another. Loss of the mitochondria-plasma membrane tethering function of the mitochondria-ER-cortex anchor (MECA) leads to a reduction in the number of ERMES contacts, accompanied by compensatory enlargement of remaining ERMES foci, revealing that the abundance and organization of one MCS can be modulated by another (Casler et al., 2025). Together, these findings highlight that MCSs are dynamically and coordinately remodeled in response to metabolic and stress conditions, enabling flexible reorganization of inter-organelle communication.

The nucleus-vacuole junction (NVJ), a MCS between the nuclear envelope and the vacuolar membrane, is formed through the interaction of the nuclear envelope protein Nvj1 and the vacuolar protein Vac8 (Jeong et al., 2017; Pan et al., 2000). Several lipid transfer and biosynthetic proteins, including Osh1, Tsc13, and Nvj2, localize to the NVJ in an Nvj1-dependent manner (Kohlwein et al., 2001; Kvam, 2004; Toulmay and Prinz, 2012). During glucose starvation (GS), the NVJ undergoes marked expansion and recruits additional factors, including the ergosterol biosynthetic enzymes Hmg1 and Hmg2, as well as regulatory proteins such as Snd3, Ypf1, Nsg1, and Nsg2, which collectively promote NVJ-dependent regulation of ergosterol biosynthesis (Fujimoto and Tamura, 2025; Hugenroth et al., 2025; Rogers et al., 2021; Tosal-Castano et al., 2021). We further revealed that the AMP-activated protein kinase Snf1, which is activated upon glucose depletion, and changes in very long-chain fatty acid (VLCFA) metabolism act upstream to control the recruitment and modification of NVJ-associated factors (Fujimoto and Tamura, 2025). The NVJ also mediates piecemeal microautophagy of the nucleus (PMN), contributing to the turnover of nuclear components under starvation conditions (Krick et al., 2008; Kvam et al., 2005; Roberts et al., 2003). Despite these advances, the molecular mechanisms that coordinate NVJ remodeling remain incompletely understood.

Msc1 (meiotic sister chromatid 1) is a poorly characterized nuclear envelope protein that has been implicated in genome maintenance. Previous work demonstrated that Msc1 reinforces homologous recombination-mediated DNA double-strand break repair during late mitosis, thereby promoting genome stability (Medina-Suárez et al., 2024). Msc1 localizes to the nuclear envelope and is required for timely completion of DNA repair prior to cytokinesis. Intriguingly, Msc1 has also been reported to exhibit a patch-like distribution along the nuclear envelope that resembles the NVJ (Breker et al., 2014; Medina-Suárez et al., 2024). Furthermore, Msc1 is stabilized by loss of Ypf1, which is a GS-dependent NVJ factor (Avci et al., 2014). These observations raise the possibility that Msc1 may function at the NVJ. However, whether Msc1 participates in NVJ remodeling or nutrient stress responses has not been investigated.

Here, we show that Msc1 is a GS-responsive NVJ factor that plays an important role in functional NVJ remodeling. We show that Msc1 expression is strongly induced during GS and that Msc1 accumulates at the NVJ in a manner dependent on the NVJ tethering proteins Nvj1 and Vac8, as well as on Snf1 signaling and VLCFA metabolism. Using biochemical and genetic analyses, we demonstrate that Msc1 localizes to the perinuclear space and acts upstream of GS-specific NVJ factors to stabilize multiple NVJ-associated proteins. In addition, we find that GS-dependent induction of *NVJ1* transcription is attenuated in *msc1Δ* cells, suggesting that proper NVJ remodeling contributes to the execution of stress-responsive transcriptional programs. Finally, we show that loss of Msc1 compromises cell survival during prolonged GS more severely than loss of the core NVJ component Nvj1. Together, these findings position Msc1 as an upstream regulator linking GS signaling to functional maturation of the NVJ and associated cellular adaptation responses.

## Results and Discussion

### Glucose starvation induces Msc1 expression and its accumulation at the NVJ

Msc1 has previously been characterized as a nuclear envelope protein that reinforces homologous recombination-mediated DNA double-strand break (DSB) repair during late mitosis, thereby contributing to genome stability (Medina-Suárez et al., 2024). Msc1 has also been reported to exhibit a patch-like localization pattern along the nuclear envelope that resembles the nucleus-vacuole junction (NVJ) (Breker et al., 2014; Medina-Suárez et al., 2024). Furthermore, The fluorescence intensity of Msc1-GFP increases upon loss of Ypf1, which is a glucose starvation (GS)-specific NVJ factor (Avci et al., 2014). These observations led us to investigate whether Msc1 functions at the NVJ during GS. To address this, we examined the intracellular localization of Msc1-GFP under different nutrient conditions. Under nutrient-rich conditions, Msc1-GFP fluorescence was barely detectable (Fig. 1A, Log). In contrast, GS, which is known to induce NVJ expansion, resulted in a dramatic increase in the Msc1-GFP signal, which prominently accumulated at the NVJ labeled with the NVJ marker Nvj1 (Fig. 1A, GS). Although NVJ expansion has also been reported during nitrogen starvation, the increase in Msc1-GFP levels under this condition was markedly weaker than that observed during GS (Fig. 1A, NS). Immunoblot analysis further confirmed that both Nvj1 and Msc1 protein levels increased modestly under nitrogen starvation but were more strongly induced under GS (Fig. 1B). These results indicate that, although NVJ expansion can be triggered by multiple nutrient stresses, Msc1 induction is preferentially coupled to GS, suggesting that Msc1 functions as a GS-responsive regulator of NVJ functions.

**Figure 1.**
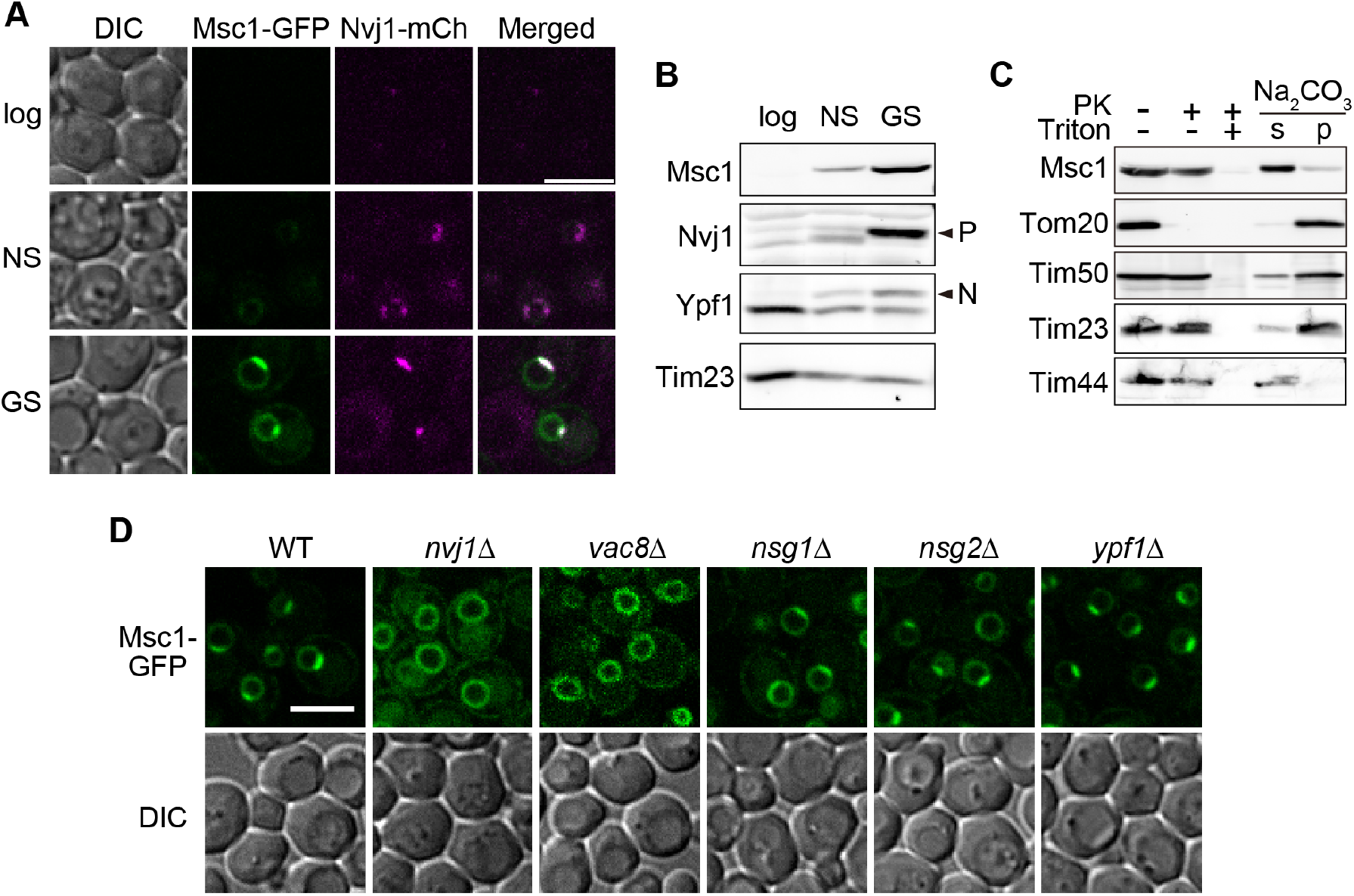
Msc1 is induced upon glucose starvation and accumulates at the NVJ. (A) Fluorescence microscopy images of cells expressing Msc1-GFP and Nvj1-mCherry under nutrient-rich conditions (Log), or after 24 h of nitrogen starvation (NS), or glucose starvation (GS). Single focal planes were shown. Scale bars, 5 µm. (B) Immunoblot analysis of whole-cell extracts prepared from wild-type cells under logarithmic growth (Log), or after 24 h of nitrogen starvation (NS), or glucose starvation (GS) conditions. Arrowheads labeled P and N represent the phosphorylated form and the stress-dependent N-glycosylated form, respectively. (C) Membrane fractions isolated from wild-type cells grown in YPLac medium were treated with 50 µg/ml proteinase K with or without 0.1% Triton X-100. Membrane fractions were treated with 0.1 M Na_2_CO_3_ and then pellets (p) and supernatants (s) were separated by ultracentrifugation. Proteins were analyzed by immunoblotting with the indicated antibodies. (D) Fluorescence microscopy images of the indicates strains expressing Msc1-GFP after 24 h of GS. Single focal planes were shown. Scale bars, 5 µm.

We also analyzed the membrane topology of Msc1. Because Msc1 is predicted to contain an N-terminal ER-targeting signal, it is expected to localize to the luminal side of the ER (Teufel et al., 2022). To test this, isolated membrane fractions were treated with proteinase K (PK). As expected, Tom20, a mitochondrial outer membrane protein with a cytosolic domain, was readily degraded. In contrast, Msc1 was resistant to PK digestion, similar to the mitochondrial inner membrane proteins Tim50, Tim23, and Tim44 (Fig. 1C). When membranes were permeabilized with Triton X-100 prior to PK treatment, all of these proteins were completely degraded (Fig. 1C). We further examined the membrane association of Msc1 by alkaline extraction. As expected, the integral membrane proteins Tom20, Tim23, and Tim50 were recovered in the pellet fraction, whereas Msc1 was efficiently released into the supernatant, similar to the peripheral membrane protein Tim44 (Fig. 1C). Together, these results indicate that Msc1 is not an integral membrane protein but instead localizes to the perinuclear space.

We next examined whether NVJ localization of Msc1 depends on known NVJ components. In cells lacking the core NVJ proteins Nvj1 or Vac8, Msc1 failed to concentrate at the NVJ and instead showed a diffuse distribution along the nuclear membrane. In contrast, deletion of GS-dependent NVJ factors, including Nsg1, Nsg2, or Ypf1, did not substantially affect the NVJ localization of Msc1-GFP. Given that Ypf1 is required for recruitment of Nsg1, Nsg2, Hmg1, and Hmg2 to the NVJ during GS (Fujimoto and Tamura, 2025), these findings suggest that Msc1 acts upstream of Ypf1 in orchestrating GS-induced NVJ functional maturation.

### Msc1 localization to the NVJ depends on Snf1 and fatty acid elongase activity

We next asked which factors are required for the GS-dependent localization of Msc1 to the NVJ. We previously showed that the AMP-activated protein kinase Snf1, which is activated upon GS, plays a central role in NVJ remodeling. We also showed that GS-induced downregulation of very long-chain fatty acid (VLCFA) elongases such as Elo3 promotes recruitment of multiple NVJ-associated factors (Fujimoto and Tamura, 2025). These findings raised the possibility that Snf1 signaling and VLCFA metabolic remodeling act upstream of Msc1. Consistent with this idea, NVJ localization of Msc1-GFP was markedly reduced in *snf1Δ* cells. (Fig. 2A, B). In contrast, deletion of *ELO3* did not significantly alter the proportion of cells exhibiting NVJ-localized Msc1-GFP but markedly expanded the size of the Msc1-positive NVJ domain (Fig. 2A, B, C, Fig. S1). Immunoblot analysis confirmed our previous findings that GS-induced phosphorylation and stabilization of Nvj1 and Nsg2, and N-glycosylation of Ypf1, are suppressed in *snf1Δ* cells but enhanced in *elo3Δ* cells (Fig. 2D). Consistent with this regulatory pattern, GS-induced accumulation of Msc1 was likewise attenuated in *snf1Δ* cells but enhanced in *elo3Δ* cells (Fig. 2D). Collectively, these results indicate that Snf1 acts upstream of Msc1 to drive GS-induced NVJ remodeling, whereas reduced Elo3 activity further accelerates this process and promotes Msc1 accumulation.

**Figure 2.**
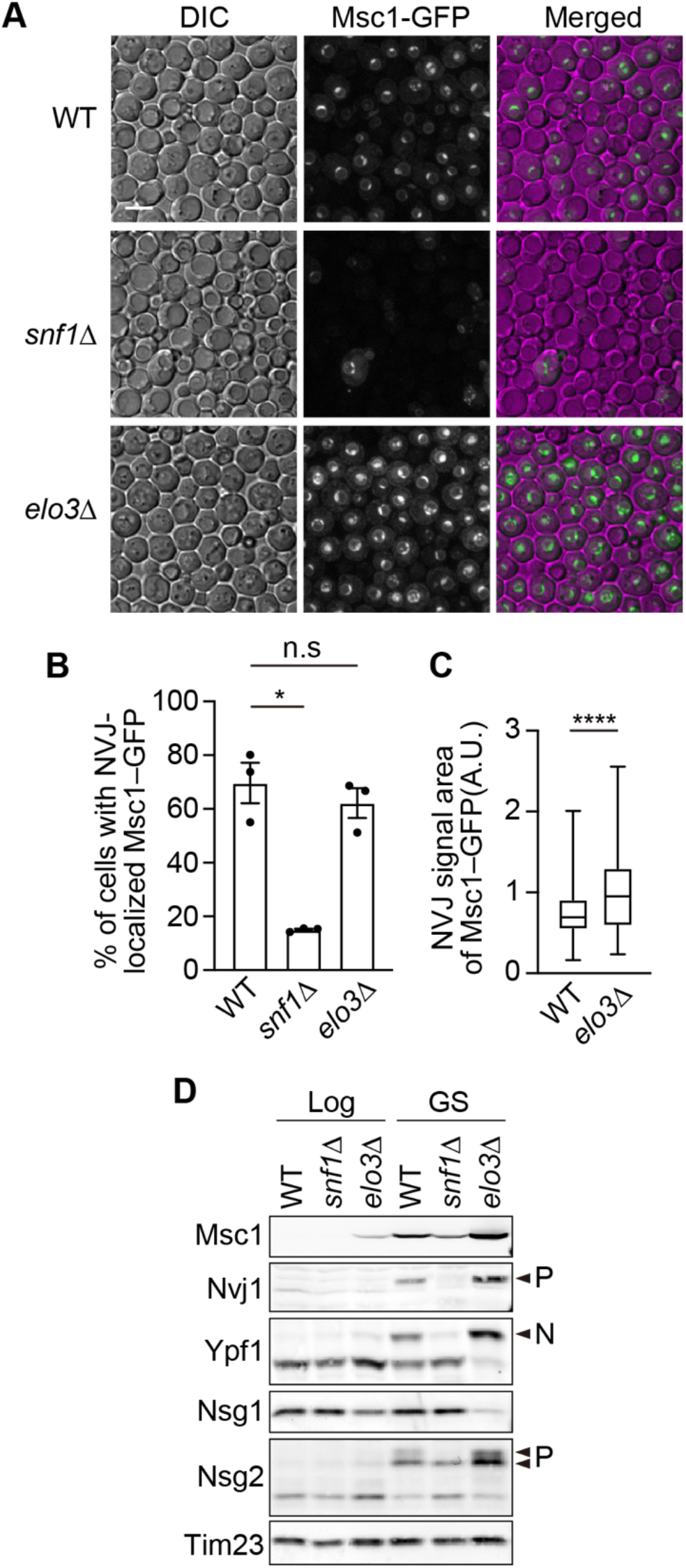
Snf1 signaling and VLCFA metabolism modulate NVJ partitioning of Msc1. (A) Representative fluorescence microscopy images of Msc1-GFP in wild-type, *snf1Δ*, and *elo3Δ* cells after 24 h of GS. Maximum projections of fluorescence images are shown. Scale bars, 5 µm. Quantification of the percentage of cells exhibiting NVJ-localized Msc1-GFP in the indicated strains after 24 h of GS. Data are shown as mean ± SEM from three independent experiments (n = 3), with 100 cells analyzed per experiment. Statistical significance was assessed using Student’s t-test; n.s., not significant, \**p* < 0.05 (C) Quantification of the size of the Msc1-positive NVJ region in wild-type and *elo3*Δ cells after 24 h of GS. Data are presented as box-and-whisker plots from three independent experiments (n = 3), with 100 cells analyzed per experiment. Statistical significance was assessed using Student’s t-test; ****P < 0.0001. (D) Immunoblot analysis of whole-cell extracts prepared from wild-type, *snf1Δ*, and *elo3Δ* cells under logarithmic growth conditions (Log) or after 24 h of GS (GS). Arrowheads labeled P and N represent the phosphorylated form(s) and the stress-dependent N-glycosylated form, respectively.

### Msc1 is required for functional remodeling of the NVJ during glucose starvation

We next sought to determine which events of GS-induced NVJ remodeling require Msc1. To examine the role of Msc1 at the NVJ under GS conditions, we generated *msc1Δ* cells and analyzed multiple hallmark events of NVJ remodeling. Deletion of *MSC1* markedly impaired GS-dependent phosphorylation and accumulation of Nvj1 and Nsg2 and caused destabilization of Nsg1 and Ypf1 (Fig. 3A). These results indicate that Msc1 plays a critical role in maintaining the stability of multiple NVJ-associated factors, including Nvj1, Ypf1, Nsg1, and Nsg2. Notably, our previous work showed that loss of Nvj1 or Ypf1 does not affect the protein levels of each other or those of other NVJ-associated factors such as Nsg1 and Nsg2 (Fujimoto and Tamura, 2025). Together with these observations, our results suggest that Msc1 plays a central role in maintaining the stability of multiple NVJ proteins during GS.

**Figure 3.**
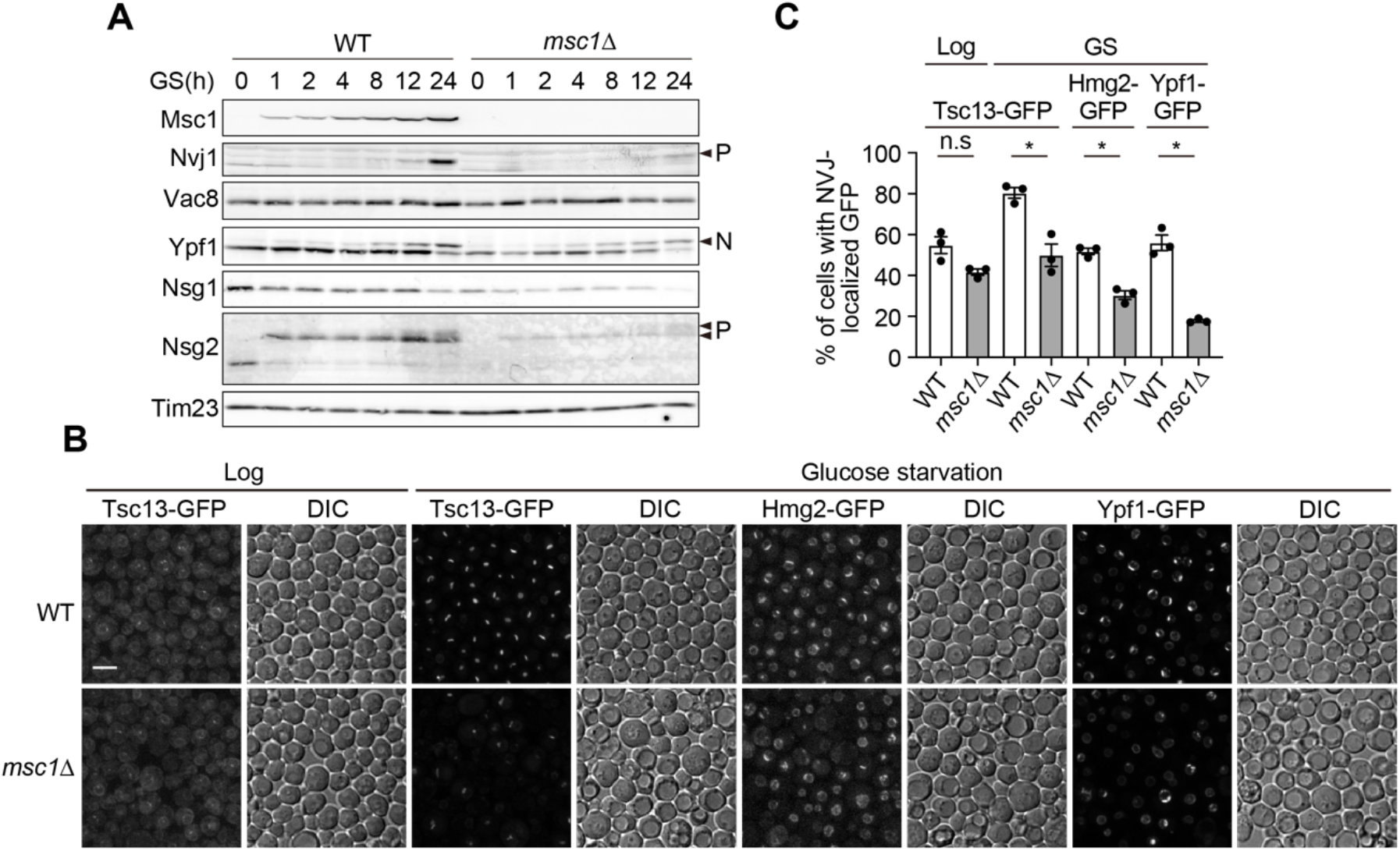
Msc1 supports stability and recruitment of NVJ-associated proteins. (A) Immunoblot analysis of whole-cell lysates prepared from wild-type and *msc1Δ* cells harvested at the indicated time points (0, 1, 2, 4, 8, 12, and 24 h) after shifting logarithmically growing cells to GS medium. Arrowheads labeled P and N represent the phosphorylated form(s) and the stress-dependent N-glycosylated form, respectively. (B) Representative fluorescence microscopy images showing localization of Tsc13-GFP, Hmg2-GFP, and Ypf1-GFP in wild-type and *msc1*Δ cells under logarithmic growth conditions (Log) or after 24 h of GS (GS). Maximum projections of fluorescence images are shown. Scale bars, 5 µm. (C) Quantification of the proportion of cells exhibiting NVJ localization of Tsc13, Hmg2, and Ypf1 following GS. Data are shown as mean ± SEM from three independent experiments (n = 3), with 100 cells analyzed per experiment. Statistical significance was assessed using Student’s t-test; n.s., not significant, \**p* < 0.05

Consistent with this notion, GS-induced recruitment of metabolic enzymes such as Tsc13 and Hmg2 to the NVJ was significantly compromised in *msc1Δ* cells. Under nutrient-rich conditions, the proportion of cells exhibiting NVJ localization of Tsc13 was comparable between wild-type and *msc1Δ* cells (Fig.3B, C). In contrast, upon GS, approximately 80% of wild-type cells displayed NVJ-localized Tsc13, whereas this fraction was reduced to ∼50% in *msc1Δ* cells. Similarly, NVJ localization of Hmg2 was observed in ∼60% of wild-type cells but in only ∼15% of *msc1Δ* cells (Fig.3B, C). These results indicate that although Msc1 is dispensable for NVJ formation, it is required for efficient GS-dependent functional maturation of the NVJ domain.

### Msc1 links NVJ remodeling to transcriptional regulation

Previous studies have shown that transcription of the *NVJ1* gene is induced upon GS (Tosal-Castano et al., 2021). To determine whether the reduced Nvj1 protein levels observed in *msc1Δ* cells under GS are due to impaired transcription or decreased protein stability, we isolated total RNA from cells in logarithmic growth phase or after GS, and quantified *NVJ1* mRNA levels by RT-qPCR. Consistent with earlier reports, *NVJ1* mRNA levels in wild-type cells increased approximately sevenfold upon GS compared with logarithmic growth conditions (Fig.4A). Notably, *MSC1* mRNA levels were dramatically upregulated (∼80-fold) following GS, suggesting that Msc1 plays a particularly important role under this stress condition (Fig.4A). In addition, *YPF1* transcription was moderately induced by GS, increasing approximately three-to fourfold (Fig. 4A). For RT-qPCR normalization, we evaluated candidate reference genes previously proposed for yeast (Teste et al., 2009) for their expression stability under GS conditions (Fig. S2). Although *ALG9* transcript levels were modestly reduced during GS, its expression remained comparable between wild-type and *msc1*Δ cells and showed the least variation among the tested candidates (Fig. S2). Based on these results, *ALG9* was used as the normalization control in subsequent analyses comparing wild-type and *msc1Δ* cells.

**Figure 4.**
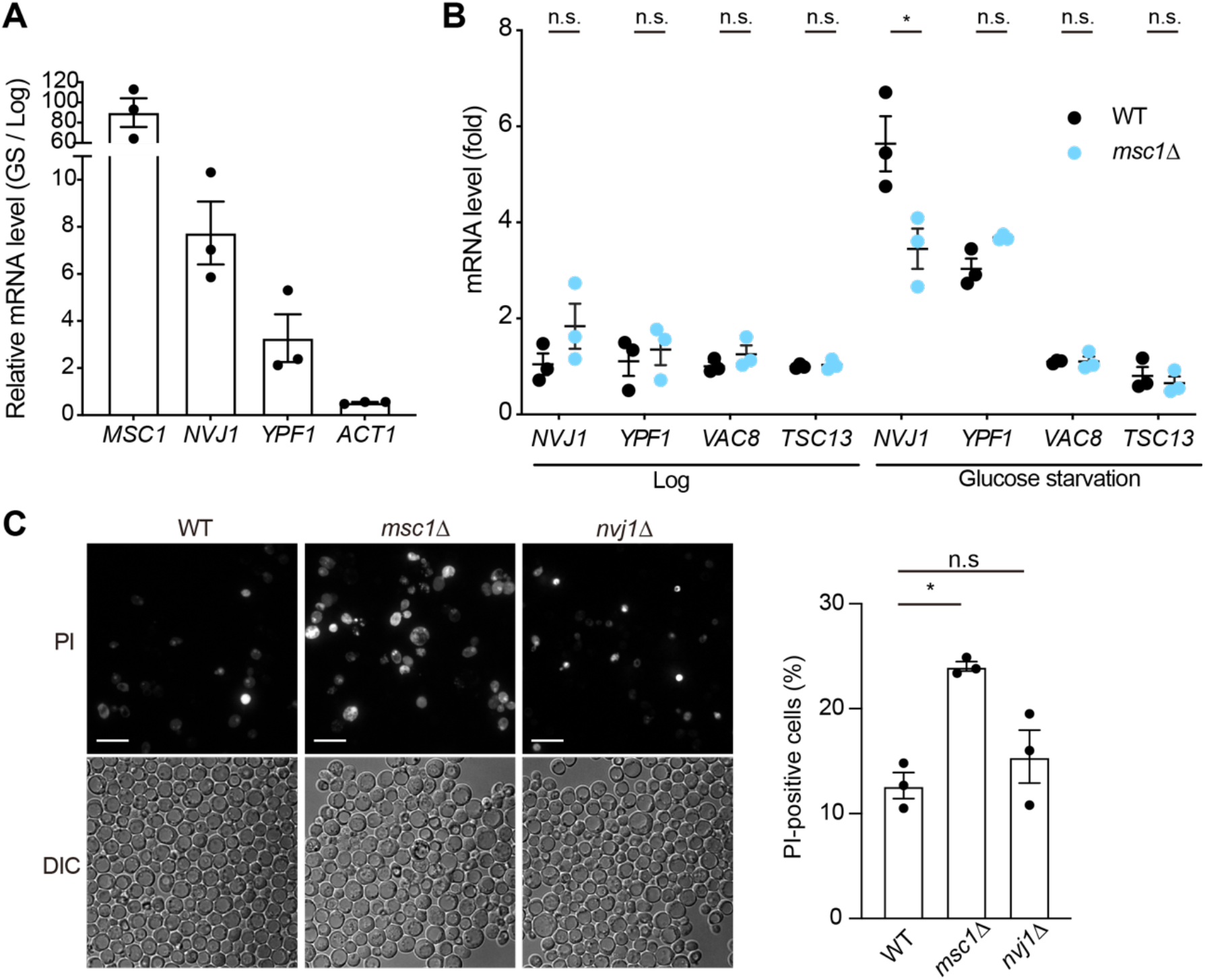
Msc1 couples NVJ remodeling to transcriptional control and stress tolerance. (A) RT-qPCR analysis of *MSC1, NVJ1, YPF1*, and *ACT1* mRNA levels in wild-type cells under logarithmic growth conditions (Log) or after 24 h of GS (GS). Transcript levels under GS conditions are shown relative to Log. Transcript levels were normalized to *ALG9*. Data represent mean ± SEM from three independent experiments. (B) RT-qPCR analysis comparing transcript levels of *NVJ1, YPF1, VAC8*, and *TSC13* between wild-type and *msc1Δ* cells under logarithmic growth conditions (Log) or after 24 h of GS (GS). Transcript levels were normalized to *ALG9*. Data are shown as mean ± SEM from three independent experiments (n = 3). Multiple unpaired Benjamin’s *t* test was applied. * *p* < 0.05. (C) Representative fluorescence microscopy images showing propidium iodide (PI) staining and corresponding DIC images of wild-type, *msc1Δ*, and *nvj1*Δ cells after 5 days of GS. Scale bar, 10 µm. Quantification of the percentage of PI-positive cells after 5 days of GS in wild-type, *msc1Δ*, and *nvj1*Δ cells. Data represent mean ± SEM from three independent experiments (n = 3), with 100 cells analyzed per experiment. Statistical significance was assessed using Student’s t-test; n.s., not significant, \**p* < 0.05

We next compared the transcript levels of NVJ-associated genes (*NVJ1, YPF1, VAC8*, and *TSC13*) between wild-type and *msc1Δ* cells. Strikingly, GS-dependent transcriptional activation of *NVJ1* was significantly suppressed in *msc1Δ* cells (Fig. 4B). In contrast, although *YPF1* transcription was modestly induced by GS, no significant difference in *YPF1* mRNA levels was observed between wild-type and *msc1Δ* cells (Fig. 4B). Transcription of *VAC8* and *TSC13* was not substantially induced by GS in wild-type and *msc1Δ* cells (Fig. 4B). Notably, several candidate reference genes (*TAF10, TFC1, UBC6*) also exhibited reduced transcript levels in *msc1Δ* cells under GS conditions, despite not being transcriptionally activated by GS in wild-type cells (Fig. S2). These observations suggest that loss of Msc1 does not cause a general defect in transcriptional activation but rather impairs the proper execution and dynamic range of GS-dependent transcriptional responses. Within this context, the robust induction of *NVJ1* appears to be particularly sensitive to Msc1 deficiency. Together, these results indicate that Msc1 contributes to transcriptional reprogramming associated with NVJ remodeling during GS. In addition to its established roles in ergosterol metabolism and nuclear microautophagy, the NVJ may therefore participate in coordinating stress-responsive gene expression programs.

Finally, we examined whether Msc1 is required for cell survival under GS. Wild-type, *msc1Δ*, and *nvj1Δ* cells were cultured in GS medium for five days, and dead cells were quantified by propidium iodide (PI) staining. Approximately 13% of wild-type cells were PI-positive under these conditions (Fig. 4C). Unexpectedly, deletion of *NVJ1* did not significantly increase cell death compared with wild-type cells (Fig. 4C). In contrast, approximately 24% of *msc1Δ* cells were PI-positive, indicating a markedly higher mortality rate than that observed in either wild-type or *nvj1Δ* cells. These results indicate that loss of Msc1 impairs NVJ function more severely than loss of Nvj1 alone. Mechanistically, deletion of *NVJ1* prevents the accumulation of Msc1 and other NVJ factors at the NVJ but does not destabilize NVJ-associated proteins. In contrast, loss of Msc1 leads to pronounced reductions in multiple NVJ proteins, including Nvj1, Ypf1, Nsg1, and Nsg2, which may contribute to the more severe functional impairment of the NVJ compared with loss of Nvj1 alone. This difference likely explains the heightened sensitivity of *msc1Δ* cells to prolonged GS.

Several important questions remain. How Msc1 is recruited to the NVJ and how it contributes to the stability of Nsg1 and Nsg2 will require further investigation. Moreover, the observation that loss of Msc1 attenuates GS-dependent induction of *NVJ1* raises the possibility that NVJ remodeling influences stress-responsive gene expression programs. Given that Msc1 localizes to the perinuclear space, it is conceivable that NVJ-associated cues may be communicated to the nucleus through yet-to-be-identified inner nuclear membrane–associated factors. Identification of Msc1-interacting proteins will therefore be an important next step toward elucidating the molecular mechanisms linking NVJ remodeling to transcriptional adaptation during glucose starvation.

## Materials and Methods

### Yeast strains and growth conditions

Saccharomyces cerevisiae strain FY833 (*MATa ura3-52 his3-Δ200 leu2-Δ1 lys2-Δ202 trp1-Δ63*) was used as background strains (Winston et al., 1995). The yeast cells used in this study are listed in Table S1. The C-terminal tagging, and gene disruptions were performed by homologous recombination using the appropriate gene cassettes amplified from the plasmids listed in Table S2. The primer pairs used in this study was summarized in Table S3.

Yeast cells were cultured in YPD (1% yeast extract, 2% polypeptone, and 2% glucose). To induce GS, YP containing 0.01% glucose was used. For nitrogen starvation, cells were cultured in SD-N medium containing 0.17% yeast nitrogen base without amino acids and ammonium sulfate, and 2% glucose. Glucose or nitrogen starvation was induced by transferring cells grown to mid-log phase in YPD medium to the respective starvation media.

### Antibodies

Polyclonal antibodies against Msc1 were generated by immunizing rabbits with N-terminally His-tagged recombinant proteins expressed in *Escherichia coli*. To generate an antigen for antibody production, a DNA fragment encoding a truncated form of Msc1 corresponding to amino acids 50 to the C-terminus (Msc1(50–last)) was amplified by PCR using yeast genomic DNA as a template and primers YU5556 and YU5557. The PCR product was digested with NdeI and XhoI and cloned into the corresponding sites of the pCold-I expression vector, resulting in an N-terminally His-tagged Msc1(50-last) construct. The resulting plasmid was transformed into BL21(DE3) cells for recombinant protein expression. Expression of His-tagged Msc1(50–last) was induced under cold-shock conditions according to the manufacturer’s instructions. The soluble recombinant protein was purified by Ni-affinity chromatography, followed by size-exclusion chromatography to remove residual contaminants. The purified protein was used as an antigen to immunize rabbits for the generation of polyclonal anti-Msc1 antibodies. The specificity of the antibody is shown in Fig. S3.

### Fluorescence microscopy

Yeast cells grown to the logarithmic phase in YPD or SCD medium, as well as cells subjected to GS, were observed by fluorescence microscopy using an IX83 inverted microscope (Olympus) equipped with a CSU-X1 spinning-disk confocal unit (Yokogawa), a 100×/1.4 NA oil-immersion objective (UPlanSApo; Olympus), and an Evolve 512 EM-CCD camera (Photometrics). Fluorescence excitation was achieved using 488-nm, or 561-nm laser lines (OBIS; Coherent) for GFP, and mCherry, respectively. Optical sections were acquired at 0.2-µm intervals throughout the entire depth of the cells, and maximum-intensity projection images were generated using ImageJ software (NIH). For staining of dead cells, propidium iodide (Merck, cat. #P4864) was added to yeast culture at a final concentration of 0.2 µg/mL from a 1.0 mg/mL stock solution, and cells were incubated at 30°C for 15 min. Cells were then collected by centrifugation, washed twice with sterile water, and resuspended for microscopic observation.

### Immunoblotting

For immunoblot analysis, whole-cell lysates were prepared from yeast cells grown to the logarithmic phase or subjected to GS. Cells corresponding to 2 OD_600_ units were collected and suspended in 300 µl of 10% (v/v) trichloroacetic acid (TCA), followed by incubation on ice for 10 min. Samples were centrifuged at 13,200 × *g* for 5 min at 4°C, and the resulting pellet was resuspended in 100 µl of 10% (v/v) TCA together with 100 µl of glass beads (0.35–0.50 mm diameter). Cells were mechanically lysed by repeated vortexing (30 s on, 60 s off, 8 cycles). After cell disruption, 900 µl of 10% (v/v) TCA was added, and samples were mixed thoroughly. Eight hundred microliters of the supernatant was transferred to a new tube and centrifuged again at 13,200 × g for 5 min at 4°C. The resulting pellet was washed with 800 µl of ice-cold acetone and centrifuged at 20,000 × g for 5 min at 4°C. The final protein pellet was dissolved in 160 µl of SDS sample buffer and incubated at 37°C for 15 min prior to SDS–PAGE. Proteins were separated using self-cast polyacrylamide gels and transferred onto PVDF membranes (Immobilon-FL; Millipore). Membranes were blocked for 1 h in blocking buffer (10 mM Tris-HCl, pH 7.5, 150 mM NaCl, 0.05% (v/v) Tween 20) containing 1% (w/v) skim milk (Morinaga, cat. #0652842). Immunoreactive signals were detected using Cy5-conjugated AffiniPure goat anti-rabbit IgG (H+L) (Jackson ImmunoResearch Laboratories) and visualized with an Amersham Typhoon scanner (Cytiva).

### Analysis of mRNA levels by quantitative RT-PCR

WT and *msc1Δ* cells were grown in YPD medium for 16 h at 30°C. Five OD_600_ unit of yeast cells was collected, and total RNA was isolated with the Direct-zol miniprep plus kit (Zyomo research, R2070). cDNA was synthesized with the PrimeScript RT reagent kit with gDNA Eraser (TaKaRa, RR047). Quantitative RT-PCR was performed with TB Green Premix ExTaq II (Tli RNaseH Plus) (TaKaRa, RR820) and the primers listed in Table S2, using Mic real-time PCR cycler (Bio Molecular Systems). Data were analyzed by the 2^−ΔΔCT^ method, normalized to the *ALG9* gene.

## Acknowledgements

We thank M. Hashimoto for her great technical assistance. We are grateful to the members of the Tamura laboratory for helpful discussion.

## Author Contributions

Conceptualization, S.F. and Y.T.; Methodology, Y.M., S.F., S.S., and Y.T.; Investigation, Y.M., S.F., S.S., and Y.T.; Writing – Original Draft, Y.T.; Writing – Review & Editing, Y.M., S.F., S.S., and Y.T.; Funding Acquisition, S.F., S.S., and Y.T.; Supervision, Y.T.

## Data and resource availability

All relevant data and details of resources can be found within the article and its supplementary information.

---

**Figure S1.**
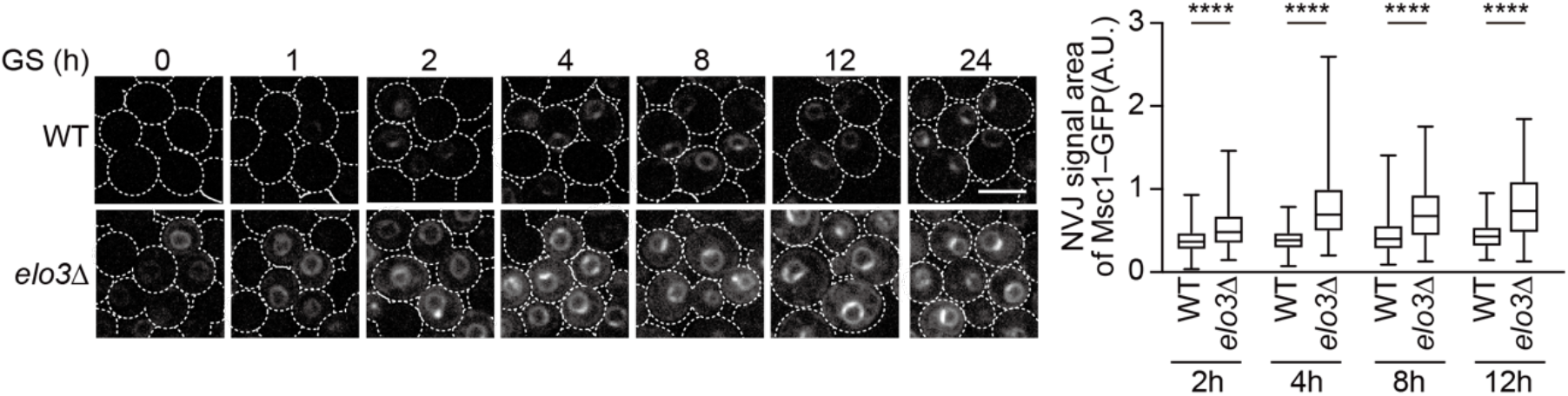
Loss of Elo3 accelerates Msc1 localization to the NVJ. Representative fluorescence microscopy images of Msc1-GFP in wild-type and *elo3Δ* cells after the indicated times of GS. Maximum projections of fluorescence images are shown. Scale bars, 5 µm. Box-and-whisker plots show the size of the Msc1-positive NVJ region in wild-type and *elo3Δ* cells after the indicated times of GS. Data are from three independent experiments (n = 3), with 100 cells analyzed per experiment. Statistical significance was assessed using Student’s t-test; \*\*\*\**p*< 0.0001.

**Figure S2.**
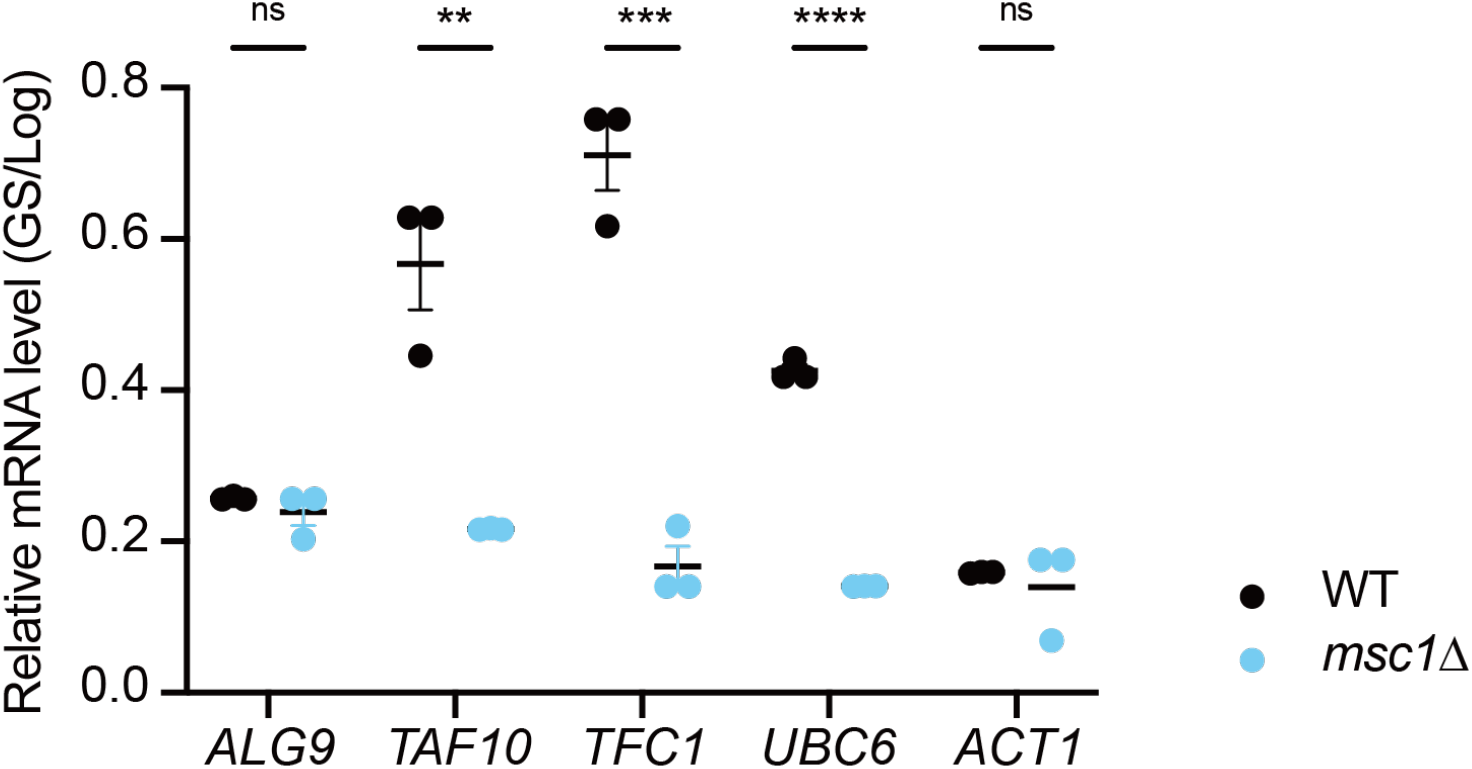
RT–qPCR analysis in log-phase and glucose-starved cells. RT-qPCR analysis of *MSC1, NVJ1, YPF1, ACT1* and *ALG9* mRNA levels in wild-type cells under logarithmic growth conditions (Log) or after 24 h of GS (GS). Transcript levels under GS conditions are shown relative to Log. Transcript levels were normalized to *TFC1*. Data are shown as mean ± SEM from three independent experiments (n = 3). Statistical significance was assessed using multiple unpaired t tests; \*\**p*< 0.01. \*\*\**p*< 0.001. \*\*\*\**p*< 0.0001.

**Figure S3.**
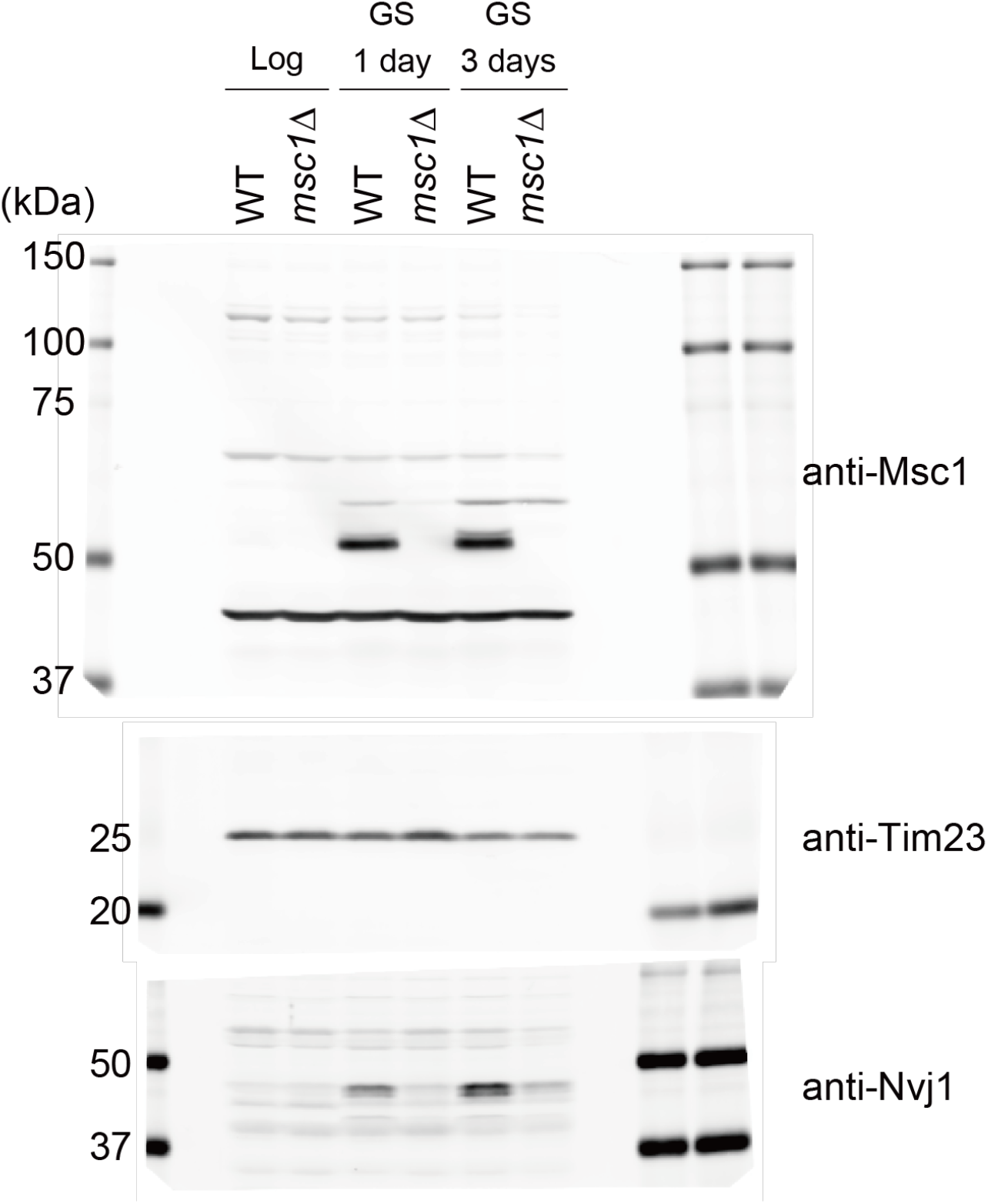
Validation of the specificity of the anti-Msc1 antibody. Total cell lysates were prepared from wild-type and *msc1Δ* cells grown to the logarithmic phase (Log) or subjected to glucose starvation (GS) for 1 or 3 days, followed by Western blotting with the indicated antibodies.

**Table S1.** Yeast strains used in this study.

| Table S1. Yeast strains used in this study. |  |  |
| --- | --- | --- |
| Name | Genotype | Identifier |
| FY833 | *MATa his3Δ200 ura3-52 leu2Δ1 lys2Δ202 trp1Δ63* | #8F5 |
| *Nvj1-mCherry Msc1-GFP* | *MATa his3Δ200 ura3-52 leu2Δ1 lys2Δ202 trp1Δ63 Nvj1-mCherry::KanMX Msc1-GFP:TRP1* | #59I5 |
| *Msc1-GFP* | *MATa his3Δ200 ura3-52 leu2Δ1 lys2Δ202 trp1Δ63 Msc1-GFP::TRP1* | #61D6 |
| *Msc1-GFP nvj1Δ* | *MATa his3Δ200 ura3-52 leu2Δ1 lys2Δ202 trp1Δ63 nvj1Δ::hphMX* | #60I7 |
| *Msc1-GFP vac8Δ* | *MATa his3Δ200 ura3-52 leu2Δ1 lys2Δ202 trp1Δ63 vac8Δ::hphMX* | #60I9 |
| *Msc1-GFP nsg1Δ* | *MATa his3Δ200 ura3-52 leu2Δ1 lys2Δ202 trp1Δ63 nsg1Δ::hphMX* | #60J3 |
| *Msc1-GFP nsg2Δ* | *MATa his3Δ200 ura3-52 leu2Δ1 lys2Δ202 trp1Δ63 nsg2Δ::hphMX* | #60J5 |
| *Msc1-GFP ypf1Δ* | *MATa his3Δ200 ura3-52 leu2Δ1 lys2Δ202 trp1Δ63 ypf1Δ::hphMX* | #60J2 |
| *Msc1-GFP snf1Δ* | *MATa his3Δ200 ura3-52 leu2Δ1 lys2Δ202 trp1Δ63 Msc1-GFP::TRP1 snf1Δ::natNT2* | #78H8 |
| *Msc1-GFPelo3Δ* | *MATa his3Δ200 ura3-52 leu2Δ1 lys2Δ202 trp1Δ63 Msc1-GFP::TRP1 elo3Δ::natNT2* | #78H10 |
| *msc1Δ* | *MATa his3Δ200 ura3-52 leu2Δ1 lys2Δ202 trp1Δ63 msc1Δ::hphMX* | #61E1 |
| *Tsc13-GFP* | *MATa his3Δ200 ura3-52 leu2Δ1 lys2Δ202 trp1Δ63 Tsc13-GFP::TRP1* | #53C7 |
| *Tsc13-GFP msc1Δ* | *MATa his3Δ200 ura3-52 leu2Δ1 lys2Δ202 trp1Δ63 Tsc13-GFP::TRP1 msc1Δ::hphMX* | #61A7 |
| *Hmg2-GFP* | *MATa his3Δ200 ura3-52 leu2Δ1 lys2Δ202 trp1Δ63 Hmg1-GFP::TRP1* | #59G2 |
| *Hmg2-GFP msc1Δ* | *MATa his3Δ200 ura3-52 leu2Δ1 lys2Δ202 trp1Δ63Hmg2-GFP::TRP1 msc1Δ::hphMX* | #61B2 |
| *nvj1Δ* | *MATa his3Δ200 ura3-52 leu2Δ1 lys2Δ202 trp1Δ63 nvj1Δ::hphMX* | #46C9 |

**Table S2.**
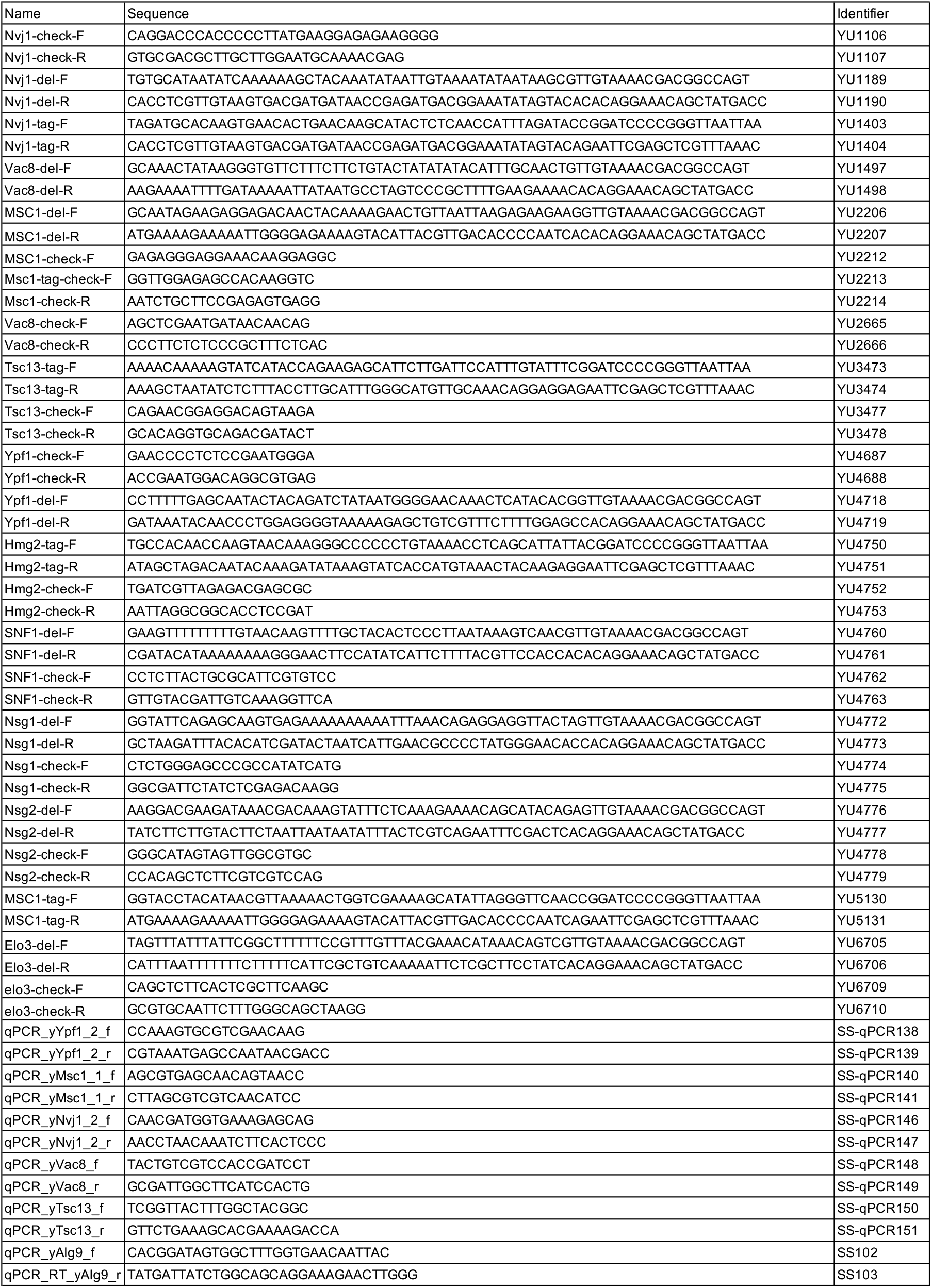
Primers used in this study.

**Table S3.** Plasmids used in this study.

| Table S3. Plasmids used in this study |  |  |
| --- | --- | --- |
| Name | Reference | Identifier |
| pBS-natNT2 | Kakimoto et al. 2018 | pYU20 |
| pBS-hphMX | Kakimoto et al. 2018 | pYU22 |
| pFA6a-GFP-S65T-TRP1 | Longtine et al., 1998 | pYU29 |
| pFA6a-mCherry-kanMX6 | Longtine et al., 1998 | pYU36 |

